# CaPTure: Calcium PeakToolbox for analysis of *in vitro* calcium imaging data

**DOI:** 10.1101/2021.09.08.458611

**Authors:** Madhavi Tippani, Elizabeth A. Pattie, Brittany A. Davis, Claudia V. Nguyen, Yanhong Wang, Srinidhi Rao Sripathy, Brady J. Maher, Keri Martinowich, Andrew E. Jaffe, Stephanie Cerceo Page

## Abstract

**Background:** Calcium imaging is a powerful technique for recording cellular activity across large populations of neurons. However, analysis methods capable of single-cell resolution in cultured neurons, especially for cultures derived from human induced pluripotent stem cells (hiPSCs), are lacking. Existing methods lack scalability to accommodate high-throughput comparisons between multiple lines, across developmental timepoints, or across pharmacological manipulations.

**Results:** We developed a scalable, automated Ca^2+^ imaging analysis pipeline called CaPTure (https://github.com/LieberInstitute/CaPTure). This method detects neurons, classifies and quantifies spontaneous activity, quantifies synchrony metrics, and generates cell- and network-specific metrics that facilitate phenotypic discovery. The method is compatible with parallel processing on computing clusters without requiring significant user input or parameter modification.

**Conclusion:** CaPTure allows for rapid assessment of neuronal activity in cultured cells at cellular resolution, rendering it amenable to high-throughput screening and phenotypic discovery. The platform can be applied to both human- and rodent-derived neurons and is compatible with many imaging systems.

## BACKGROUND

Transient increases in intracellular calcium levels are a requisite for neuronal activity, thus providing a useful strategy for measuring cellular activity at both network-wide and single-cell resolution. Calcium imaging is frequently used to measure activity dynamics in the brains of awake, behaving animals using fiber photometry or miniaturized microscopes coupled with endoscopic imaging [1–3]. This field is advancing rapidly with the advent of new technologies [4], and a number of computational methods have been developed to analyze calcium imaging data *in vivo* at both the single-cell level as well as to assess bulk calcium dynamics within the entire field of view [5–8].

However, due to differences in signal-to-noise ratios and background fluorescence in intact tissue versus cell culture systems, collecting and analyzing calcium imaging data from *in vitro* cell culture models requires different computational approaches [9]. For example, *in vitro* cell model systems are comparatively less active and more synchronous than intact brain samples. Many of the existing methods for calcium imaging analysis detect changes in activity, and then combine those synchronous signals into the signal attributed to a single cell [10]. However, due to the high degree of synchronicity in *in vitro* systems, these methods erroneously combine activity measurements for multiple cells that are firing as an ensemble. With advancements in human induced pluripotent stem cell (hiPSC) technologies and *in vitro* genetic modelling of disease, the need to accurately measure neuronal activity in cultured neurons has become increasingly important. As current models often involve either co-culture systems with multiple species as source material (e.g. rodent glial cells co-cultured with human neurons) or mixed cell-type assemblages (e.g. primary cortical tissue, or hiPSC-derived organoids), genetically encoded calcium indicators (GECIs) enable important cell-type specific targeting. Thus, strategies for measuring neuronal activity that use AM-dye based Ca^2+^ indicators or multi-electrode arrays, where a priori targeting or characterization of a specific cell population is not feasible, result in limited cell-type specific information.

Acquisition of this information enables comparisons between hiPSC lines derived from different individual donors, or from transgenic rodent models. In review of existing literature, we found most analysis methods require a high degree of user input to define parameters [11, 12], or extensive knowledge of the data being acquired to provide information for specific functions, like *‘findpeaks’* in MATLAB. On the other hand, FluoroSNNAP - Fluorescence Single Neuron and Network Analysis Package - accurately detects events, but is GUI based. Hence this package is not compatible with high performance computing clusters, and, in our hands, the pre-processing failed to accurately remove noise from the data [13]. Utilizing a field-based thresholding approach requires a high degree of similarity between all acquired time-lapse movies, or the selection of amplitude and intensity thresholds to be performed for each field independently.

Here we introduce CaPTure, which is an automated analysis pipeline that facilitates 1) the accurate detection of neurons, 2) the identification of calcium events in individual cells, and 3) the calculation of image-based network connectivity metrics. Utilizing a GECI that co-expresses a fluorescent marker protein in the cell type of interest, we extended the FluoroSNNAP software package by introducing additional data pre-processing steps to detect regions of interest (ROIs) to focus subsequent analysis, and normalize fluorescence intensity over time [13]. We added data-driven motifs representing events observed in our data, and calculated synchrony metrics including clusters of synchronous cells to assess ensemble activity. Compared to existing analysis methods, our method accurately quantifies dynamic measurements in selected cells, while incorporating both per-field and individual per-ROI neuronal activity metrics. Thus, this method has the advantage of facilitating comparisons of neuronal and network activity between genetic models of disease and pharmacological manipulations. The method is highly amenable to parallel computing and high-throughput screening.

## IMPLEMENTATION

### Sample preparation

#### hiPSC-derived neurons

Fibroblast donors were male and of European ancestry - these research subjects were enrolled in the Sibling Study of Schizophrenia at the National Institute of Mental Health in the Clinical Brain Disorders Branch (NIMH, protocol 95M0150, NCT00001486, Annual Report number: ZIA MH002942053, DRW PI) as previously described [14]. Early passage fibroblasts (<5 passages) were reprogrammed into hiPSCs as previously described [15], and subsequently differentiated through neural progenitor stages into cortical neurons. Neurons were co-cultured in 24-well ibidi plates with astrocytes prepared from the cortices of neonatal rats to promote neuronal maturity as previously described [14, 16]; and were maintained with partial media changes twice a week for up to 10 weeks (Day in Vitro (DIV70)).

#### Animals

Wild-type mice were bred for the generation of postnatal day 0 mice primary neuronal cultures. Mice were purchased from Jackson laboratories (Bar Harbor, ME, C57BL6/J; stock #000664). Timed-pregnant Wistar rats for astrocyte cultures were obtained from Charles River Laboratories (Wilmington, MA, USA; stock Crl:WI003). Rodents were housed in a temperature-controlled environment with a 12:12 light/dark cycle and *ad libitum* access to standard laboratory chow and water. All experimental animal procedures were approved by the SoBran Biosciences Institutional Animal Care and Use Committee.

#### Mouse primary cortical cultures

Mouse cortical neurons were cultured on a 24-well ibidi plate (Cat. No. 82406, ibidi GmbH, Munich, Germany) as previously described with modifications [17]. Briefly, on the day of birth, mice were anesthetized by being placed on ice, then rapidly decapitated and their cortices removed. Cortical tissue was dissociated using papain, and plated at a density of 2.5×10^5 per well on a 24-well ibidi plate coated with poly-D-lysine and laminin. Neurons were maintained in culture with partial media changes every two days and imaged between DIV14 and DIV15.

#### Viral Transduction

hiPSC-derived neurons were transduced at DIV23 with adeno-associated virus expressing mRuby2 and GCaMP6s under the control of a synapsin promoter (MOI ∼6×10^4, Addgene viral prep # 50942-AAV1 [18]. Following a full media exchange on DIV25, neurons were cultured for at least 21 days and imaged on DIV 42 or 63. Mouse primary cultures were transduced with 1:10 viral concentration used in human experiments of the same virus (human synapsin 1 promoter was ubiquitously expressed in mouse neurons). Mouse primary cultures were infected at DIV5-DIV8 prior to DIV14-DIV15 recordings.

### Imaging Acquisition

#### LSM780 confocal microscope

Primary mouse cortical cultures and hiPSC-derived neurons were imaged in culture media on a Zeiss LSM780 equipped with a 10X/0.45NA objective, a temperature- and atmospheric-controlled enclosure to maintain neurons at 37° and 5% CO2. A reference image was acquired for each field of mRuby fluorescence followed by a time-series was acquired at 4Hz for 8 minutes. In some cases, tetrodotoxin (TTX, 1uM) was then added to block synaptic transmission and incubated for at least 5 minutes prior to imaging to equilibrate.

#### Spinning Disk confocal microscope

Neurons were removed from culture media and were continuously perfused with artificial cerebro-spinal fluid (ACSF) containing (in mM): 128 NaCl, 30 glucose, 25 HEPES, 5 KCl, 2 CaCl_2_, and 1 MgCl_2_ (pH 7.3) [16]. Imaging was performed at DIV56 or DIV70 on a custom-built Zeiss Spinning Disk confocal with a 20X/1.0NA water immersion objective. A reference image was acquired using mRuby fluorescence, then a time-series was acquired at 10Hz for 5 minutes. For experiments in which pharmacological blockers were added, TTX (1uM) was included in the perfusate for at least 5 minutes prior to imaging.

#### Acquisition parameters

From all scopes, two image types are collected: a time-series of GCaMP6s fluorescence and a reference image of mRuby to demarcate infected neurons. The reference image of the LSM780 scope is downsampled using the MATLAB function *imresize* to match the time-series image in X and Y dimensions.

| Scope | Reference image<br>X Y Z | Pixel to<br>micron | Time-series image<br>X Y Z | Pixel to micron |
| --- | --- | --- | --- | --- |
| LSM 780 | 1024x1024 pixel<br>850.19x850.19 μm | 0.83x0.83 μm<br>per pixel | 256x256 pixel<br>850.19x850.19 μm | 3.32x3.32 μm<br>per pixel |
| Spinning disk | 1024x640 pixel<br>660.48x412.8 μm | 0.645x0.645<br>μm per pixel | 1024x640 pixel<br>660.48x412.8 μm | 0.645x0.645 μm<br>per pixel |

#### Toolbox installation and software requirements

All data processing for CaPTure is conducted in MATLAB (Version 2017a or later). The processing pipeline is divided into several steps as described below, the execution of which are explained in the following repository https://github.com/LieberInstitute/CaImg_cellcultures. The repository consists of a ‘toolbox’ directory whose path needs to be added to the MATLAB working directory to run any of the processing steps. The directions to download and install the toolbox are described in the *‘installation’* step of the repository.

#### Statistics

To calculate the effect of pharmacological manipulations, the lmerTest R package was used for performing linear mixed effects modeling as a function of treatment main effect (Baseline versus TTX) and used cell line and the cell culture experimenter as the random intercepts.

### Using CaPTure

In this section we describe the analysis workflow: first we identify ROIs by segmenting neurons in the cell-fill channel, and then extract fluorescence intensity. Then we identify “peaks,” which are used to calculate per-image and per-cell summary and aggregate metrics to assess network and cellular activity.

#### Step1: Convert .czi time series files to .mat files

A time-series of images was collected for each imaging field, and saved using the Zeiss proprietary .czi file format to maintain image metadata. Since all of the image data processing is performed in MATLAB, we recommend that users convert the raw data to MATLAB format for fast and easy access. We use the Bio-Formats package called ‘bfmatlab’ ([Linkert et al. 2010](https://sciwheel.com/work/signin?targetUrl=%2Fwork%2F%23%2Fitems%2F1399038%2Fdetail%3Fcollection%3D451597)) to load the .*czi* data into MATLAB and use traditional save functions in MATLAB to save to .*mat* format. The ‘bfmatlab’ package supports the conversion of multiple proprietary file formats obtained from different microscope systems, thus enabling the use of CaPTure on calcium imaging data obtained from various systems.

#### Step2: Identify ROIs

CaPTure allows the user to automate detection of ROIs, and then to select ROIs based on their shape or size. The strategy allows us to detect cells that express the cell-type specific GECI, but are inactive. From each reference image, we identify infected neurons from which to measure calcium dynamics (**Figure 1A**). Neurons have a complex morphology, and we aimed to identify signal from the soma, and not from surrounding neuropil. Thus, we used the MATLAB function *‘imhmin’* to suppress the background signal coming from the neurites (**Figure 1B**). We then used the *region growing* technique ([Kroon 2008](https://sciwheel.com/work/signin?targetUrl=%2Fwork%2F%23%2Fitems%2F11082085%2Fdetail%3Fcollection%3D451597)) for segmenting ROIs from the red image, where the pixel with the minimum fluorescence intensity of the image is chosen as the initial seed location, and the region is iteratively grown by comparing all unallocated neighboring pixels to the seed region. The difference between the intensity value of each pixel and the mean of the region is used as a measure of similarity. The pixel with the smallest difference measured this way is allocated to the respective region. This process stops when the intensity difference between the region mean and that of the new pixel becomes larger than a user specified threshold, in this case, the standard deviation of the image (**Figure 1C**). The fully grown region is termed the background, thus leaving out the regions with high intensity which become the final segmented ROIs (**Figure 1D**). To select for neurons and to remove noise, debris and neuropil from further inclusion in the data, we used eccentricity (a measure of the roundness of the ROI calculated by the MATLAB function *‘regionprops3’*) and a minimum size threshold to filter out ROIs from neuropil and noise (**Figure 1E, F**). The output of Step 2 provides the identification of all ROIs.

**Figure 1.**
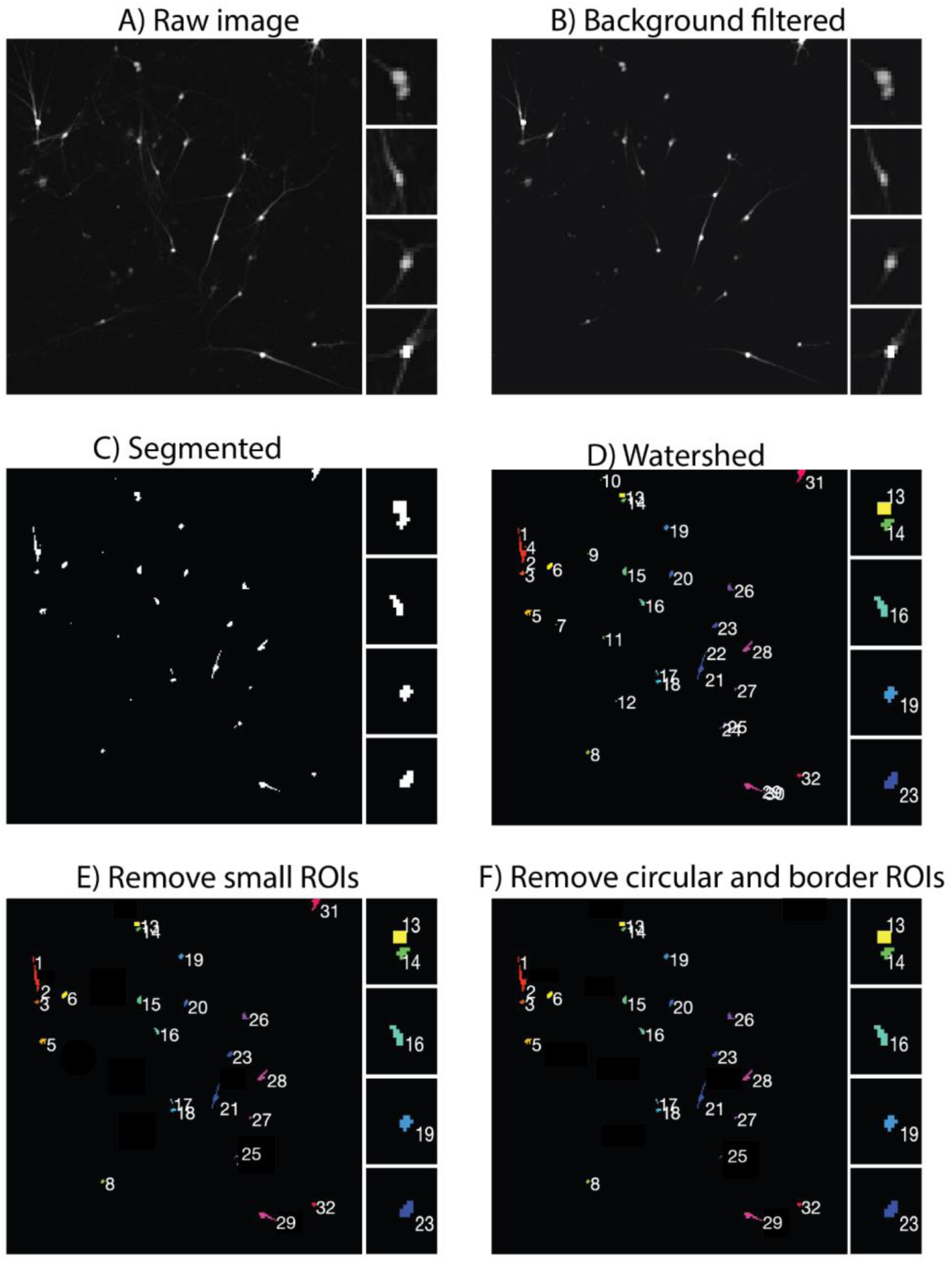
Nuclei Segmentation: **A**) Raw *‘.czi’* image of nuclei cultures. **B**) Background filtered image of the raw *‘.czi’* image. We used the MATLAB function *‘imhmin’* to suppress the background noise coming from the neurites. **C**) Segmented (using *‘Region Growing’*) binary image of nuclei. **D**) Final watershed segmentation of ROIs from the binary image (inserts of ROI 13 and 14). Watershed was performed for better extraction of individual ROIs that are spatially in close proximity. **E**) ROIs (e.g., 4,7,11) that are smaller in size (total pixels in the segmented region) are excluded from the final segmentation. **F**) ROIs with an eccentricity >0.99 and ROIs (i.e., 31) on the image border are excluded.

#### Step3: Extract traces from each ROI

Calcium imaging allows for measurement of calcium levels in each individual cell by measuring dynamic fluorescence intensity. From each identified neuron, i.e., ROI, we extract calcium signals by measuring the fluorescence intensity over time. Traces (signal) are extracted from the green video using the ROI segmentations from Step 2. Each point on the trace is the average intensity of all the pixels of the segmented ROI at that Z frame in the green video. The output of Step 3 (**Figure 2)** is raw traces for each ROI. For ease of illustration, in subsequent figures we focus on three ROIs: ROI 16-low activity (light teal), ROI 19-moderate activity (medium blue) and ROI 23-high activity (royal blue).

**Figure 2.**
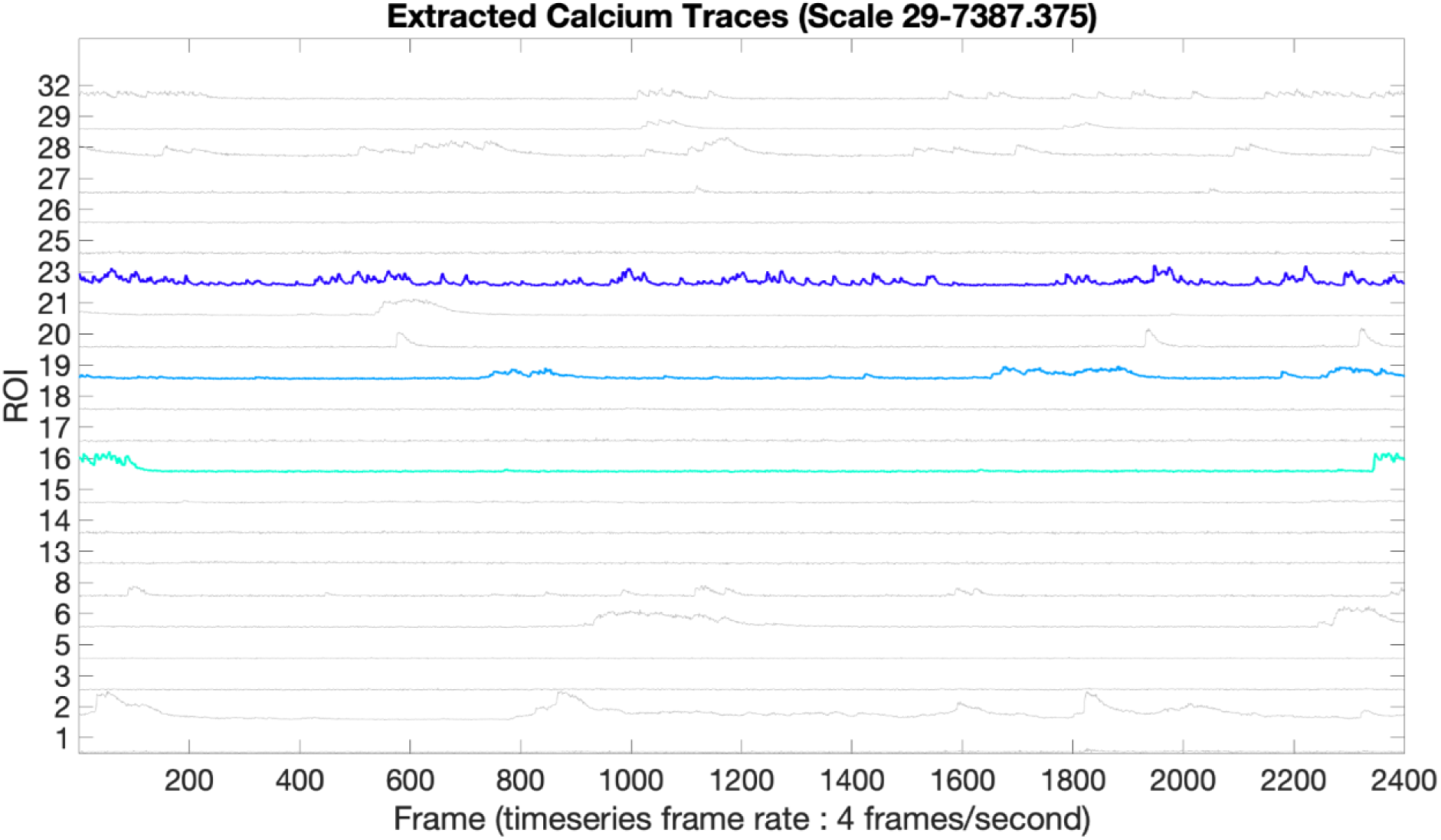
Extracted Calcium Traces: The graph shows the calcium activity for each ROI segmented in Step2, and highlights traces 16, 19, and 23 as examples of low-, medium- and high-activity ROIs, respectively. The x-axis is the frame number of the time series and the y-axis is the ID for the segmented ROI. These traces have a minimum of 29 and the maximum of 7387 units of mean fluorescence intensity.

#### Step4: Extract delta fluorescence/fluorescence (dff) from step3

Dynamic fluorescent intensity is normalized to baseline. Due to fluctuations in viral transfection efficiency, baseline activity, expression of the virus and the position of the cell within the sample, there can be differences in the baseline fluorescence intensity between ROIs. Thus, we normalize each trace using a rolling average as described in Developing And Assessing Methods For Calcium Imaging Data (2.2.4 Smoothing: DFF) ([Jia et al. 2011](https://sciwheel.com/work/signin?targetUrl=%2Fwork%2F%23%2Fitems%2F28490%2Fdetail%3Fcollection%3D451597)). The output of Step 4 provides normalized traces with smoothing (**Figure 3**).

**Figure 3.**
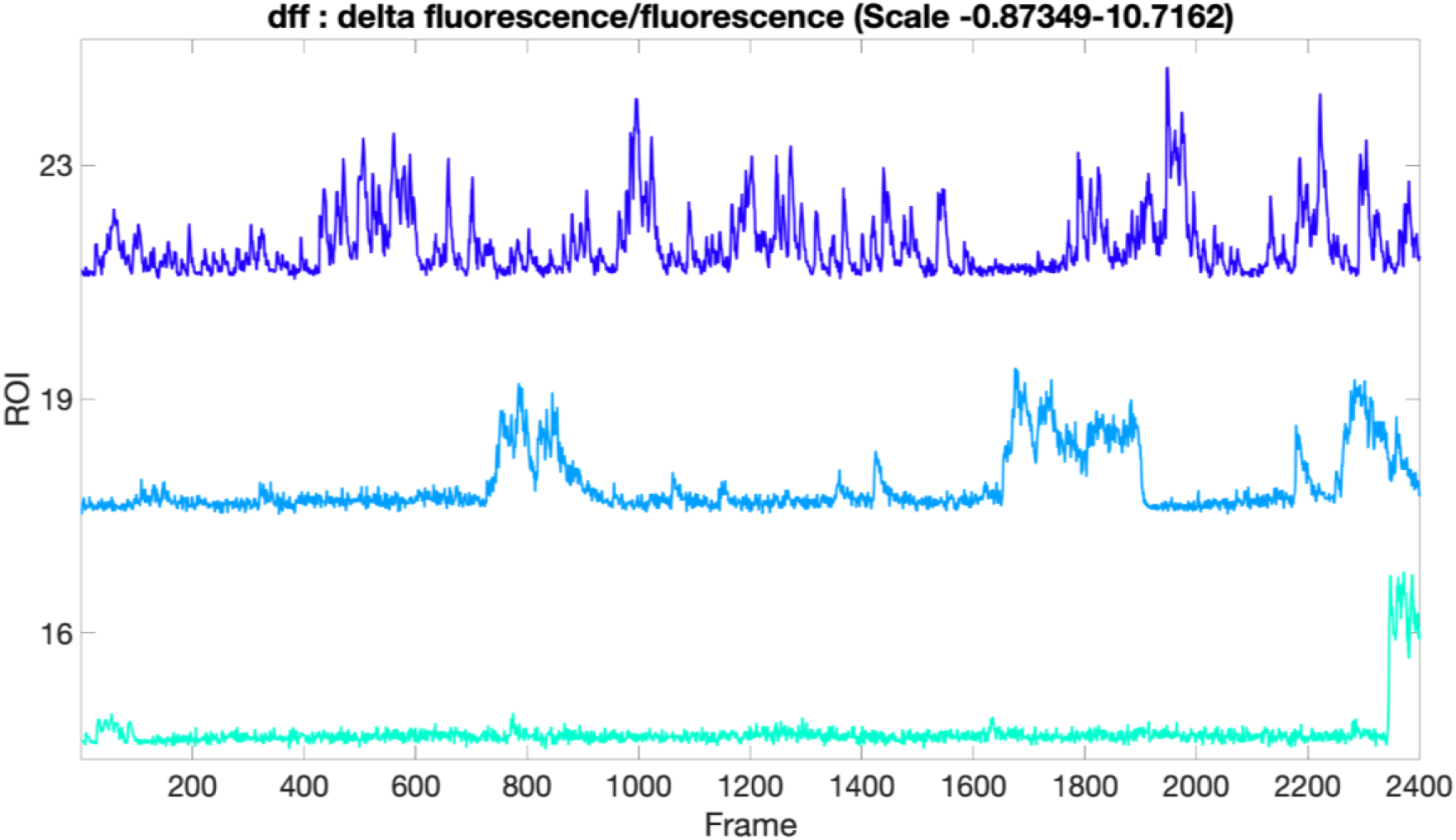
Delta fluorescence/fluorescence: The graph shows the normalized calcium traces extracted from the calcium activity shown in Figure 2. The x-axis is the frame number of the time series and the y-axis shows only segmented ROIs 16, 19, 23 (low, medium and high activity) for easy visualization.

#### Step5: Construction of correlation maps

To identify calcium events, a correlation map is constructed to compare the pattern of fluorescence intensity changes with known motifs representing calcium events. Prior to the calculation of the correlation map, the dff traces needed to be interpolated because the motif library, created by FluoroSNNAP ([Patel et al. 2015](https://sciwheel.com/work/signin?targetUrl=%2Fwork%2F%23%2Fitems%2F955401%2Fdetail%3Fcollection%3D451597)), utilized a frame rate of 10 frames/second (**Figure 4A**). We utilized the FluoroSNNAP motifs and constructed seven motifs based on observations from our data (**Figure S1**). A matrix (‘Ca’, rows = motifs, columns = x axis of the trace) of correlation coefficients of all motifs across the trace is computed (**Figure 4B**). The correlation coefficients are set to a value of zero at locations across the trace where the intensity/height of the trace are below a certain threshold that represents the background, to avoid noise (**Figure 4C**). The output of Step 5 aligns normalized traces to motifs (**Figure 4**).

**Figure 4.**
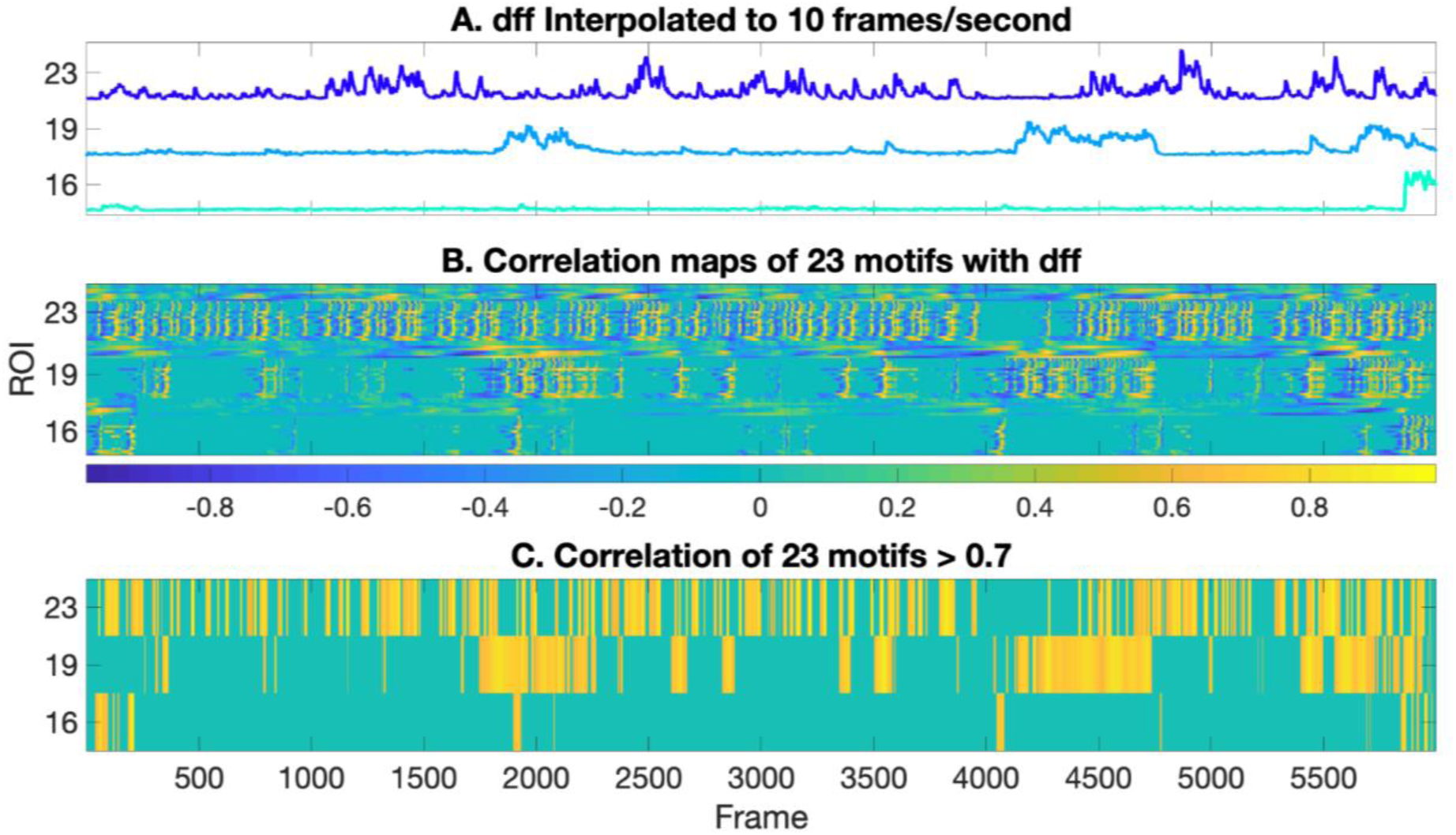
Motif Correlation Maps: A) The normalized traces (4 frames/second) from step4 are interpolated to 10 frames/second to match the frame rate of the motifs being correlated. B) Motif correlation map showing frames in yellow when the event predominantly matches a motif, and frames in blue when the event is least matched with the same motif. The frames in turquoise represent the background. C) Showing only frames where the maximum correlation (of 23 motifs) is above a threshold of 0.7.

#### Step6: Extract event location and duration

We next extract the event location and duration for each event in each ROI (**Figure 5)**. A final row matrix is computed by picking the maximum correlation coefficient from each column of ‘Ca’. The points that exceed the user given correlation threshold (0-1) on the row matrix represent the events of that trace. A high correlation threshold might result in missing some events, while a low correlation threshold will potentially pick noise as events, so an optimal threshold of ∼0.7 was used for our datasets (**Figure 5B**). The total number of all the consecutive points/frames that cross the threshold is taken as the event duration in frames. The output of Step 6 counts and classifies motifs (**Figure 5**). We illustrate the occurrence of each motif in our example data set (**Figure 6A**), and the occurrence of each motif within each ROI (**Figure 6B**).

**Figure 5.**
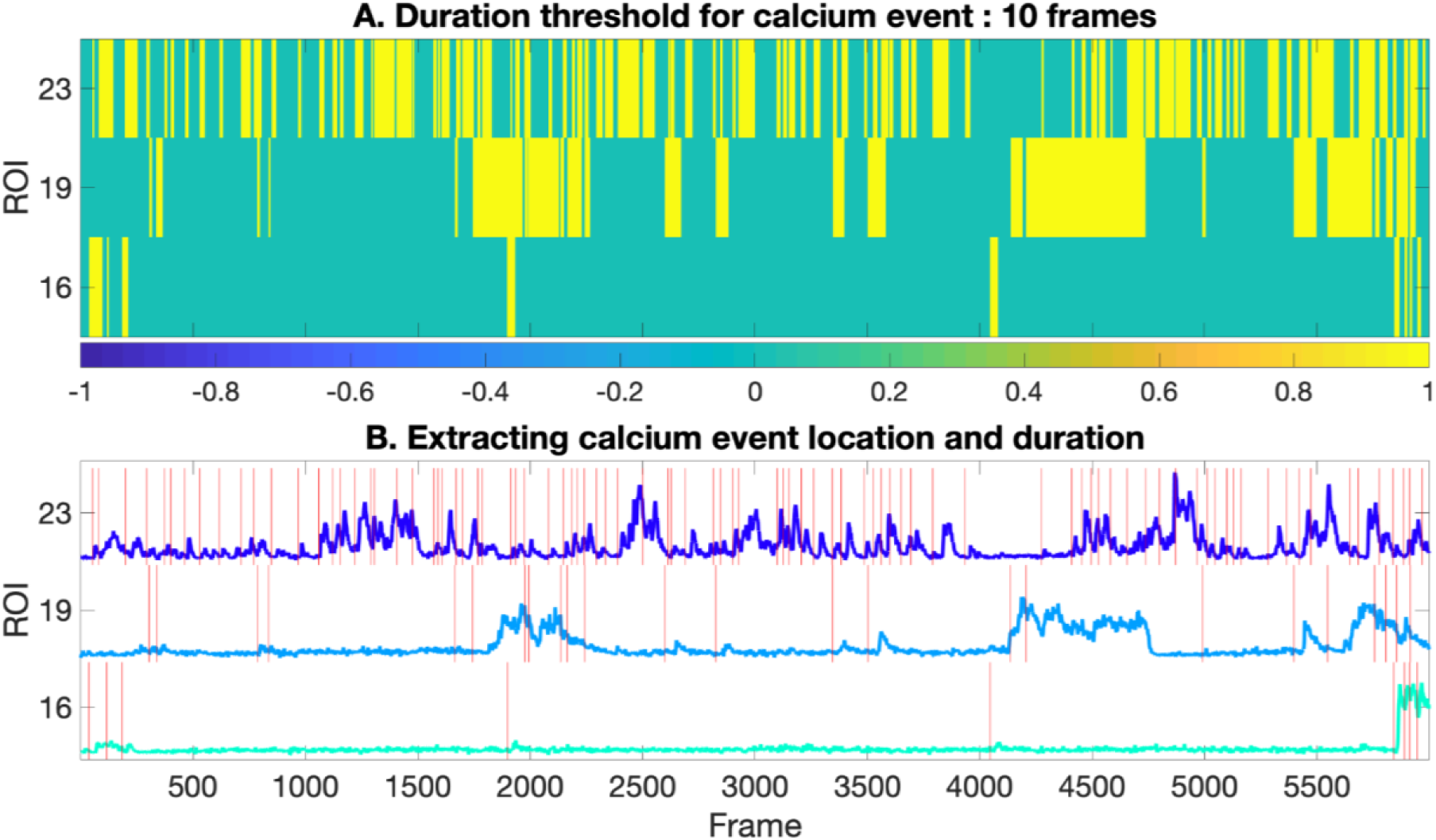
Extract calcium events: **A**) The thresholded correlation map from Figure 4 is converted to a binary map. **B**) Extracted event location and duration based on the binary map in (**A**).

**Figure 6.**
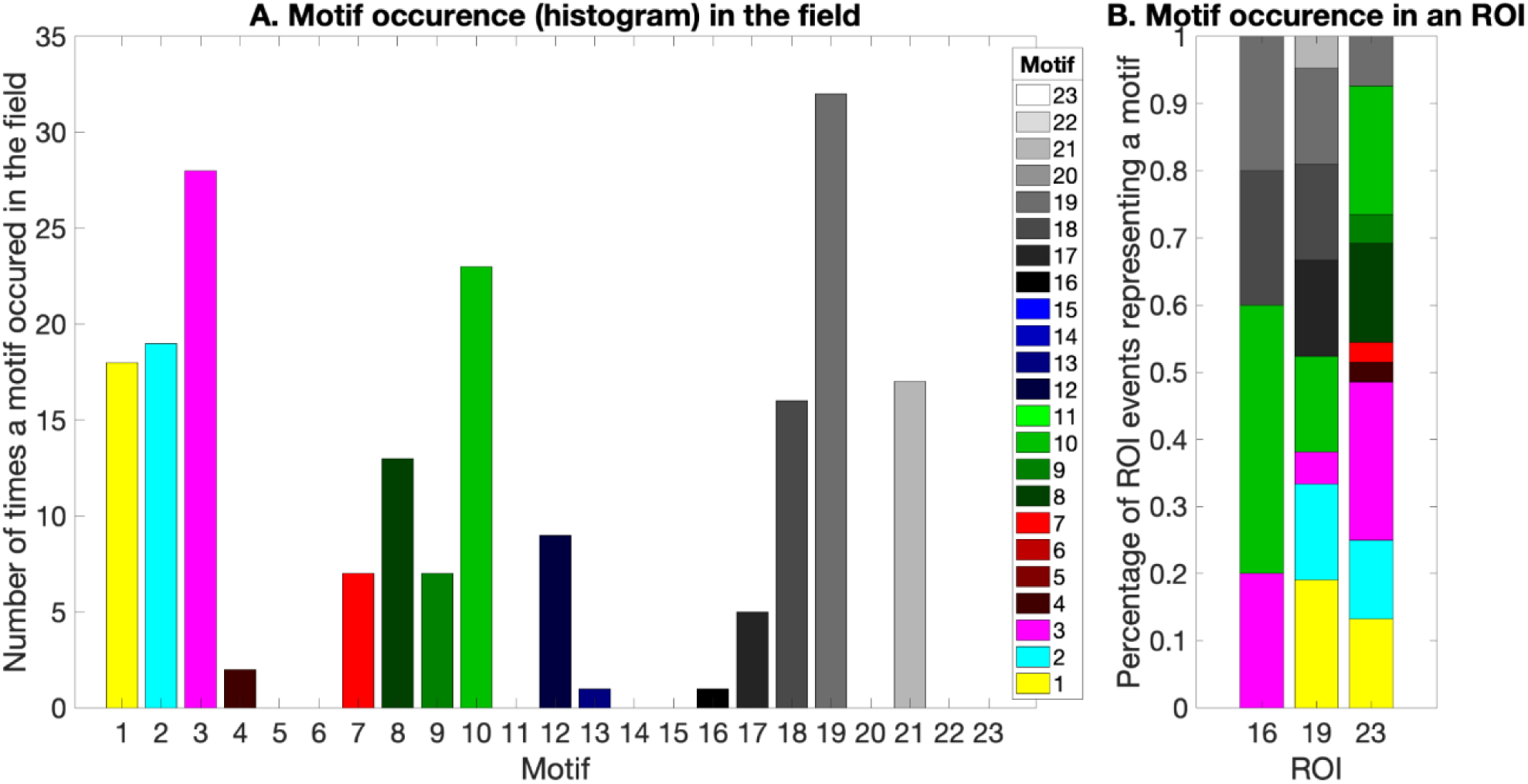
Frequency of a motif occurrence: **A**) The barplot shows the frequency of occurrence of a motif in the specific field. The x-axis shows individual motif and the y-axis shows the total number of times the motif appeared in the field. **B**) The barplot shows the percentage of events in a ROI that correlates with a specific motif. The x-axis shows the ROIs 16, 19, 23 and the y-axis shows the percentage of events of the ROI.

#### Step 6A (optional): Synchronicity

Because neurons in *in vitro* networks are highly interconnected, we aimed to estimate the degree to which calcium events were synchronous across a given field. To do this we quantified how synchronous the calcium activity is between the ROIs of a given field using the functions (*‘SCA’*) provided by the FluoroSNNAP package (**Figure 7**). The package provides different methods to quantify synchrony including phase correlation, entropy, and Fourier Transforms of the calcium traces and events. We used the correlation method applied on calcium activity and corresponding surrogate traces of pairwise neurons in a field to quantify the network synchronicity [13]. The output of Step 6 shows the degree to which events in each ROI are correlated with events in other ROIs.

**Figure 7.**
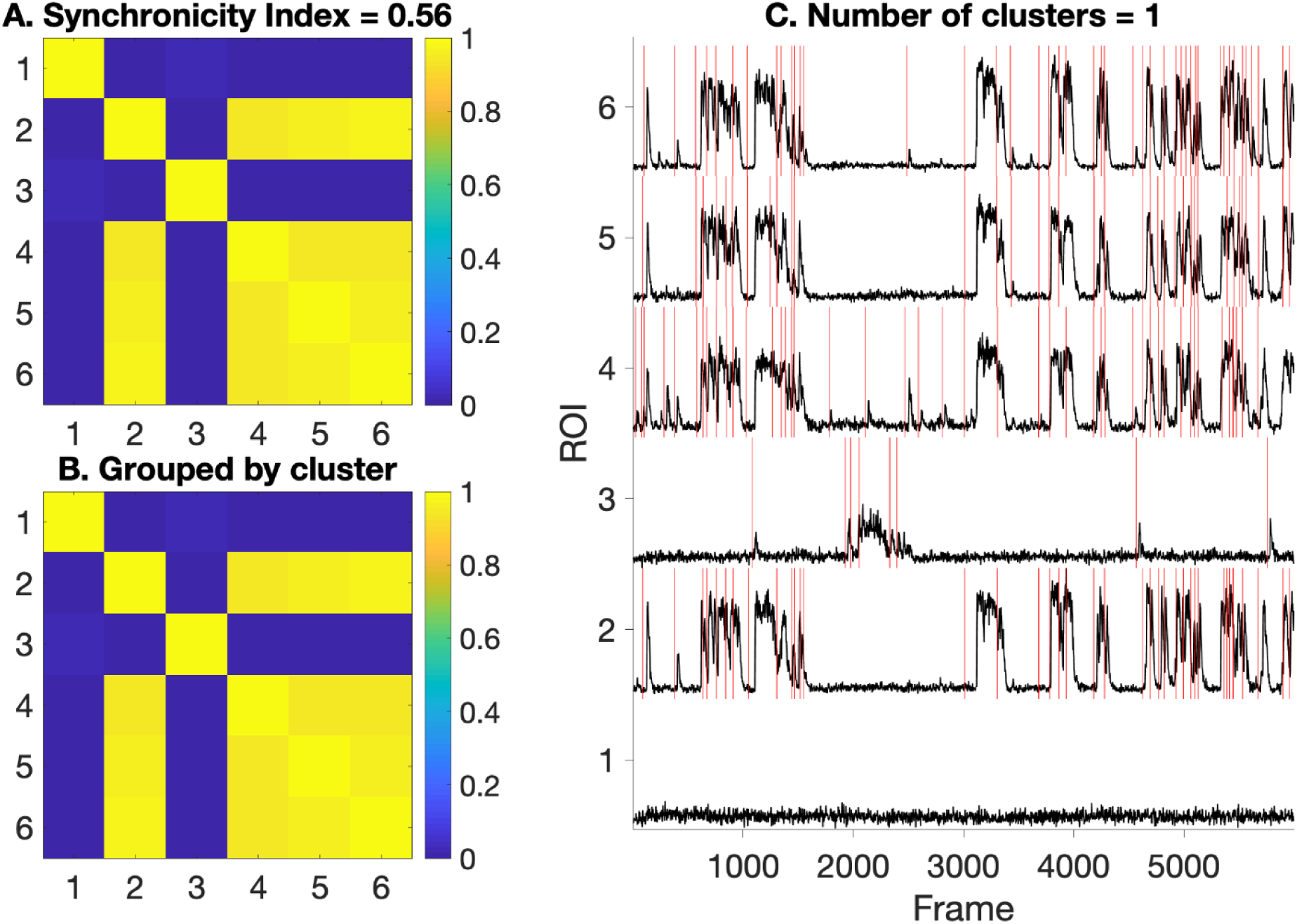
Network Synchronicity: **A**) Heatmap showing pairwise correlation of the calcium activity in the field. Synchronicity Index (0-1) represents a measure for network synchrony of the field. **B**) Correlation map regrouped by the identified synchrony clusters. **C**) Showing calcium activity of the field to visually analyze network synchronicity.

#### Step7: Extract Final Data

A custom MATLAB script was written to extract two types of metrics: individual ROI metrics in the file *long_dat* and image metrics in the file *man*. This allows us to make comparisons across individual cells and across fields. The final *man.csv* file represents the image level summary statistics (in columns) for each image (in rows) in the dataset.

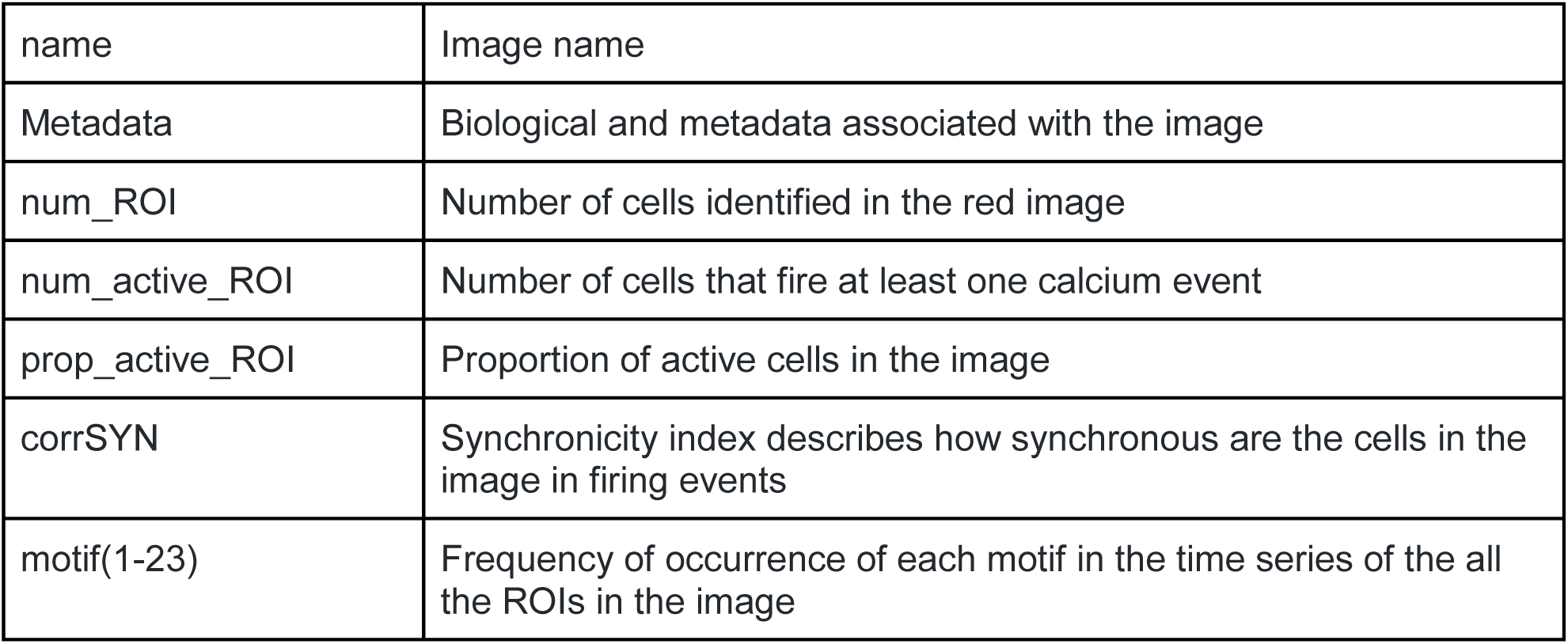

|  |  |
| --- | --- |
| name | Image name |
| Metadata | Biological and metadata associated with the image |
| num_ROI | Number of cells identified in the red image |
| num_active_ROI | Number of cells that fire at least one calcium event |
| prop_active_ROI | Proportion of active cells in the image |
| corrSYN | Synchronicity index describes how synchronous are the cells in the image in firing events |
| motif(1-23) | Frequency of occurrence of each motif in the time series of the all the ROIs in the image |

The final *long_data.csv* file represents the ROI level summary statistics (in columns) for each ROI (in row) in the dataset.

|  |  |
| --- | --- |
| name | Image name which the ROI belongs to |
| Metadata | Biological and metadata associated with the image |
| events_ROI | Number of calcium events that a cell produced |
| avg_width | Average duration (frames) of events for that ROI |
| Volume | Number of pixels in red image that corresponds to the ROI |
| Eccentricity | Describes if the ROI is more elongated or more circular in shape. An ROI whose eccentricity is 0 is actually a circle, while an ROI whose eccentricity is 1 is a line segment. |
| motif(1-23) | Frequency of occurrence of each motif in the time series of the ROI |

## RESULTS

To demonstrate the utility of the workflow, we apply CaPTure to several *in vitro* preparations of neurons (e.g mouse primary cortical neurons and hiPSC-derived neurons), and demonstrate versatility by applying the workflow to data acquired on an additional microscope system. Finally, we demonstrate the robustness of the method by blocking neuronal activity in iPSC-derived neurons with pharmacological agents and assessing algorithm performance.

We first applied this toolbox to mouse cortical neurons in culture. These cultures are both more dense and more mature than hiPSC-derived neuronal cultures. We confirmed that our ROI detection method accurately identified ROIs and extracted calcium events in an active, dense mouse culture system (**Table 1, Figure 8A, 8B**). We identified unique patterns of synchronicity, which suggests that some sets of neurons preferentially fire together (**Figure 8C**). We then extract image- and ROI-based metrics for final data analysis (**Figure 8D**).

| Sample | Cells | ROIs | Active ROIs | Avg field activity/ 60 secs | Synchronicity | Clusters |
| --- | --- | --- | --- | --- | --- | --- |
| Figure 1 | Human | 22 | 16 | 0.9 events/ROI | 0.03 | 4 |
| Figure 8 | Mouse | 41 | 41 | 3.48 events/ROI | 0.08 | 3 |

**Figure 8.**
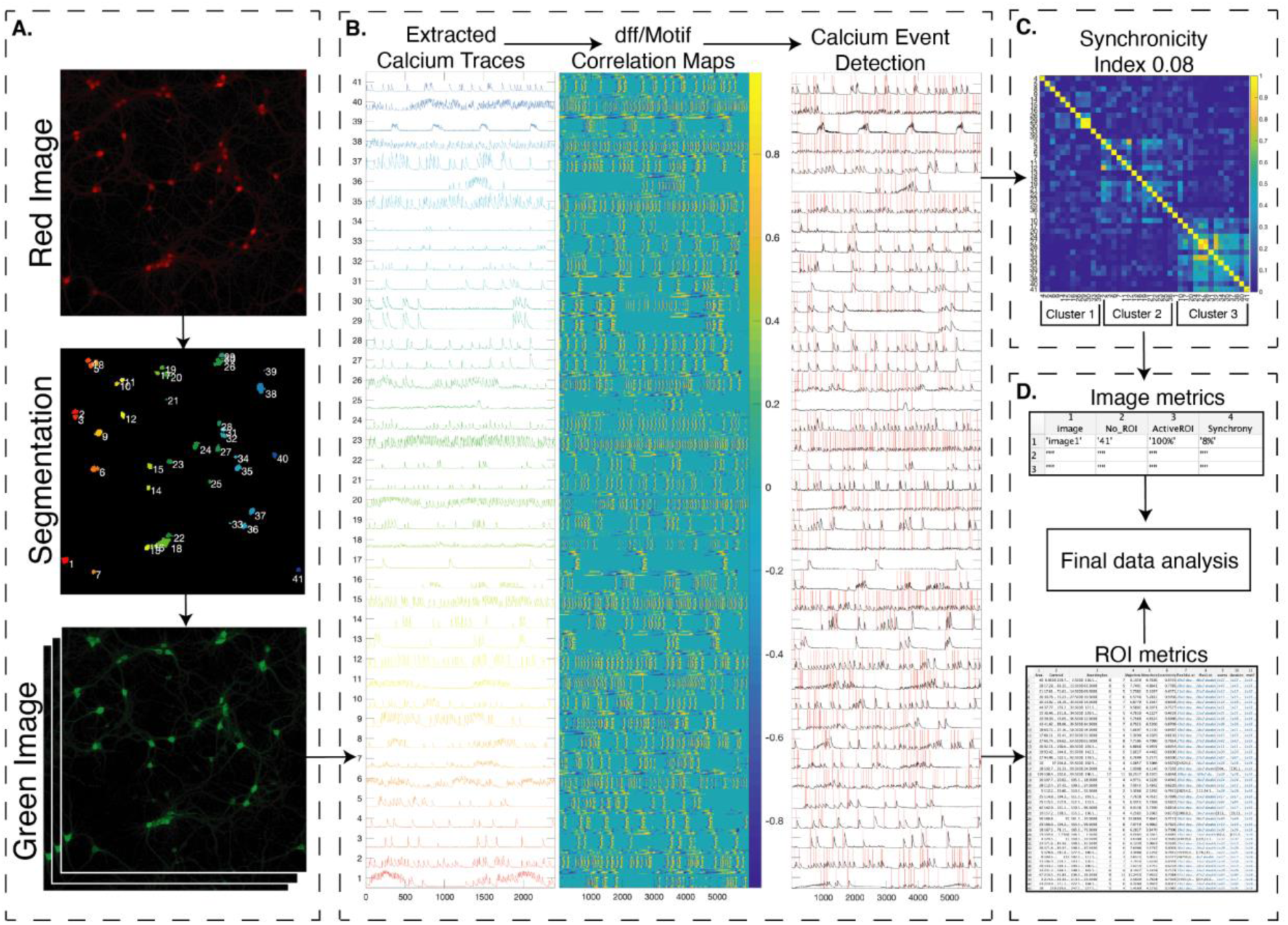
Primary mouse cortical neuronal culture data processed through CaPTure: **A**) The raw red ‘.czi’ image of mRuby-expressing neurons, its corresponding color coded neuronal segmentation and the GCaMP6s time-series. **B**) Extracted raw calcium traces, the motif correlation map of the interpolated dff traces and event detection on the interpolated dff traces using the motif correlation maps. **C**) Correlation map of inter-neuron calcium activity grouped by the cluster. **D**) Final extracted metrics at image(field) level and ROI(neuron) level.

Additionally, we applied CaPTure to images acquired on a higher-resolution microscope with a smaller field of view. An additional challenge with this data set was the presence of physical drift of the sample due to the continuous perfusion of ACSF over the coverslip containing neurons. Thus, time-lapse images needed to be registered such that movement in the X and Y directions would be computationally removed. We therefore registered the images by aligning each frame iteratively to the preceding frame, and then aligning the green GCaMP6s time-lapse to the red cell-fill image (**Figure 9A**, [Catallini 2020](https://sciwheel.com/work/signin?targetUrl=%2Fwork%2F%23%2Fitems%2F11315540%2Fdetail%3Fcollection%3D451597)). To demonstrate the utility of this approach, we show the correlation of the fluorescence of each frame with the mean of the entire timeseries (**Figure 9B**). Prior to registration, the mean of the timeseries is not highly correlated, and the correlation is variable by frame. After registration, correlation of each frame is highly correlated. Following registration, we applied CaPTure to this set of data, identifying ROIs and individual peaks (**Figure 10)**.

**Figure 9.**
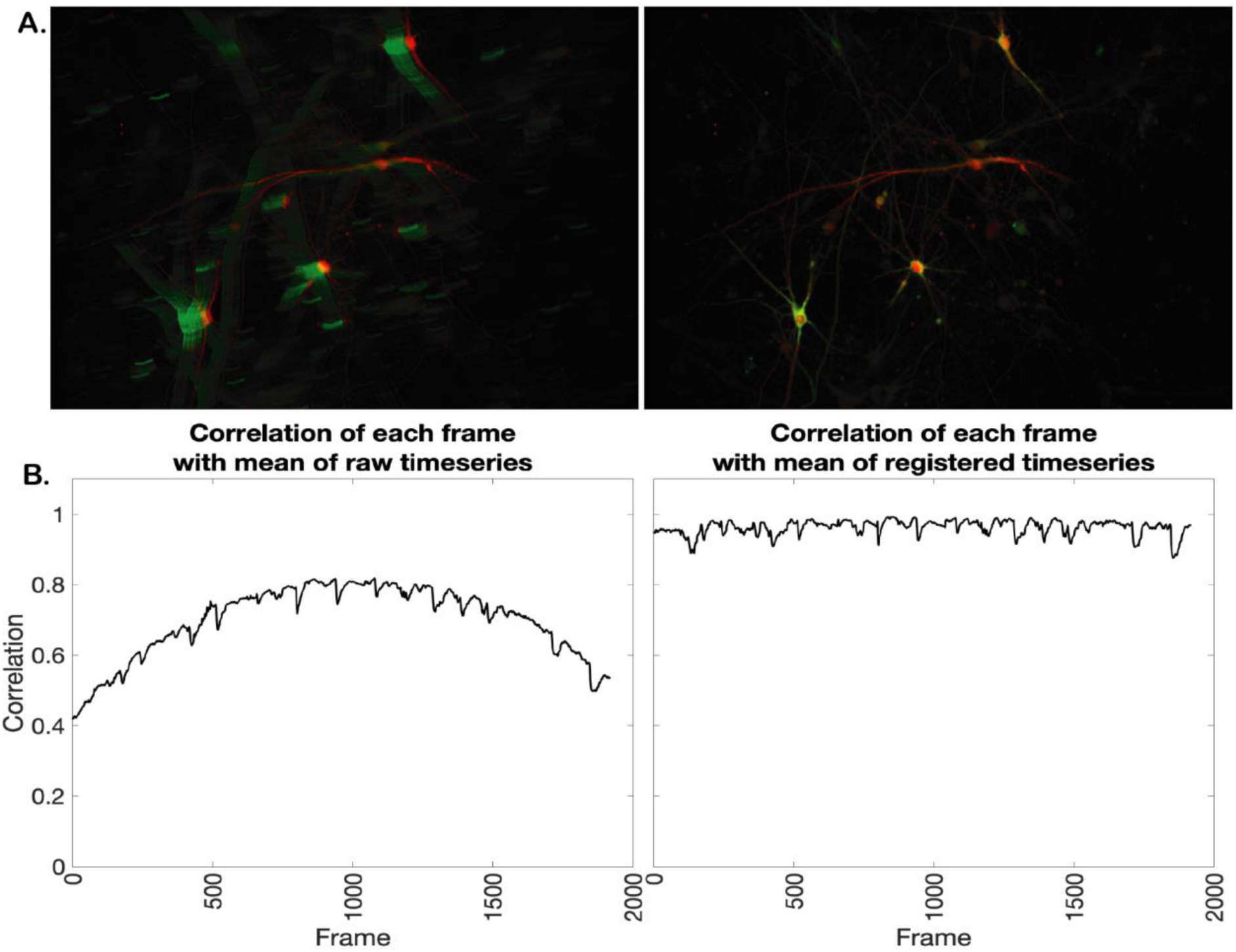
Image Registration to correct physical drift: **A**) Red image overlaid on the maximum intensity projection of the green time series of the raw data (left) and the registered data (right). **B**) Graphs showing correlation (y-axis) of each frame (x-axis) with the mean intensity image of the time series for raw data (left) and registered data (right).

Finally, we tested the accuracy of our peak detection methodology by applying tetrodotoxin (TTX), a pharmacological agent that blocks sodium channels, thus preventing neuronal activity. We measured calcium transients in neurons before and after the application of TTX (**Figure 11A, B**). In this case, the background intensity or the height threshold used in building Motif correlation maps are estimated based on the baseline, not based on the total data including the manipulation (**Figure 11C**). We see a decrease in the number of calcium events per ROI following TTX treatment (**Figure 11D**; mean ± SEM from 7 lines, Baseline 13.83 ± 0.349; TTX 2.84 ± 0.0683, linear mixed effects model p-value <2e-16). This demonstrates that CaPTure is accurately detecting synaptic events.

**Figure 11.**
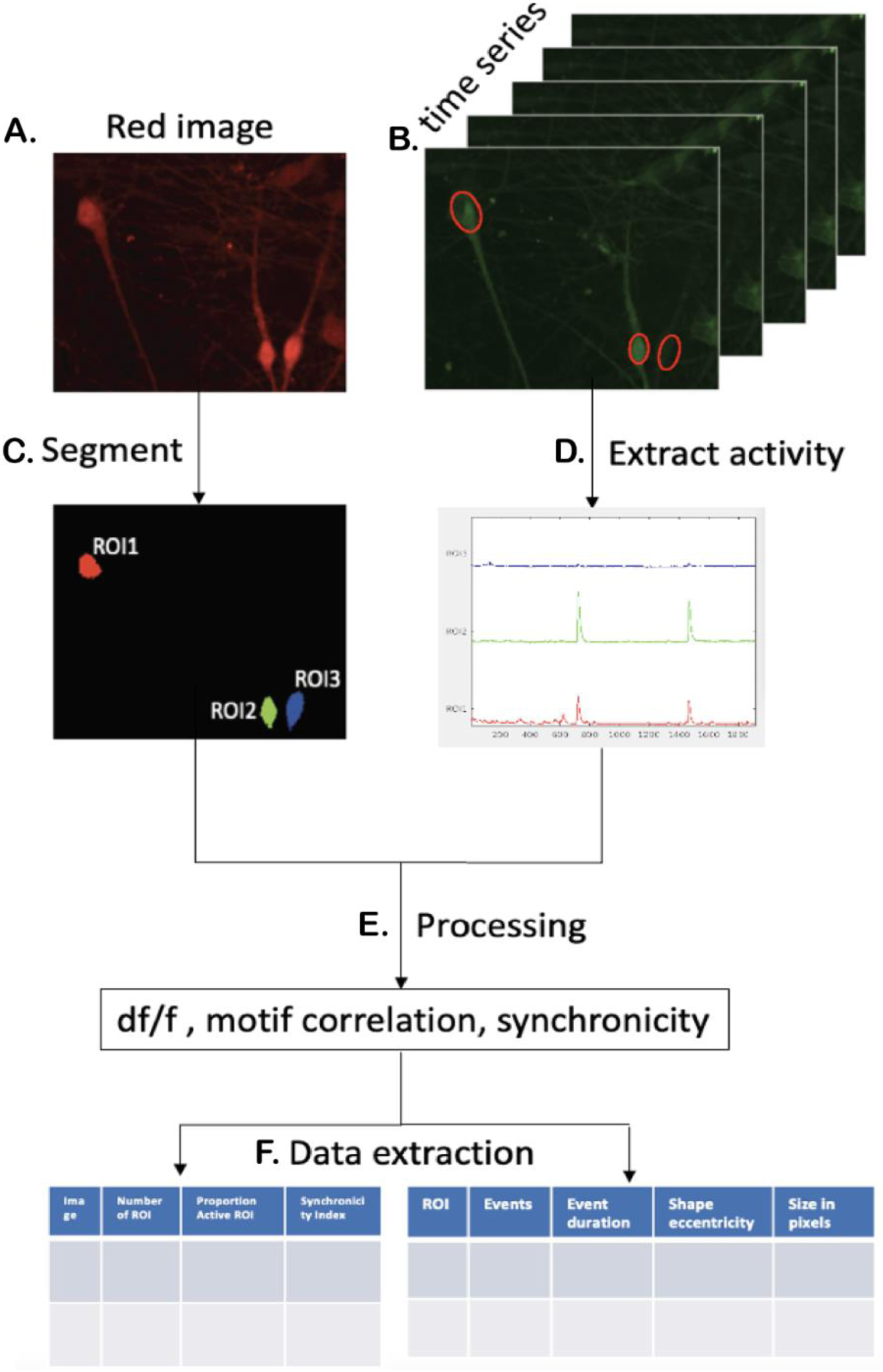
CaPTure applied to secondary data set: **A**) Raw *‘.czi’* image of nuclei and **B**) the corresponding raw *‘.czi’* time series. **C**) Segmentation of nuclei image. **D**) Calcium activity extracted from times series for each segmented nuclei. **E**) Different processing steps used in CaPTure. **F**) Final metrics extracted into tables for image level and ROI level data.

**Figure 12.**
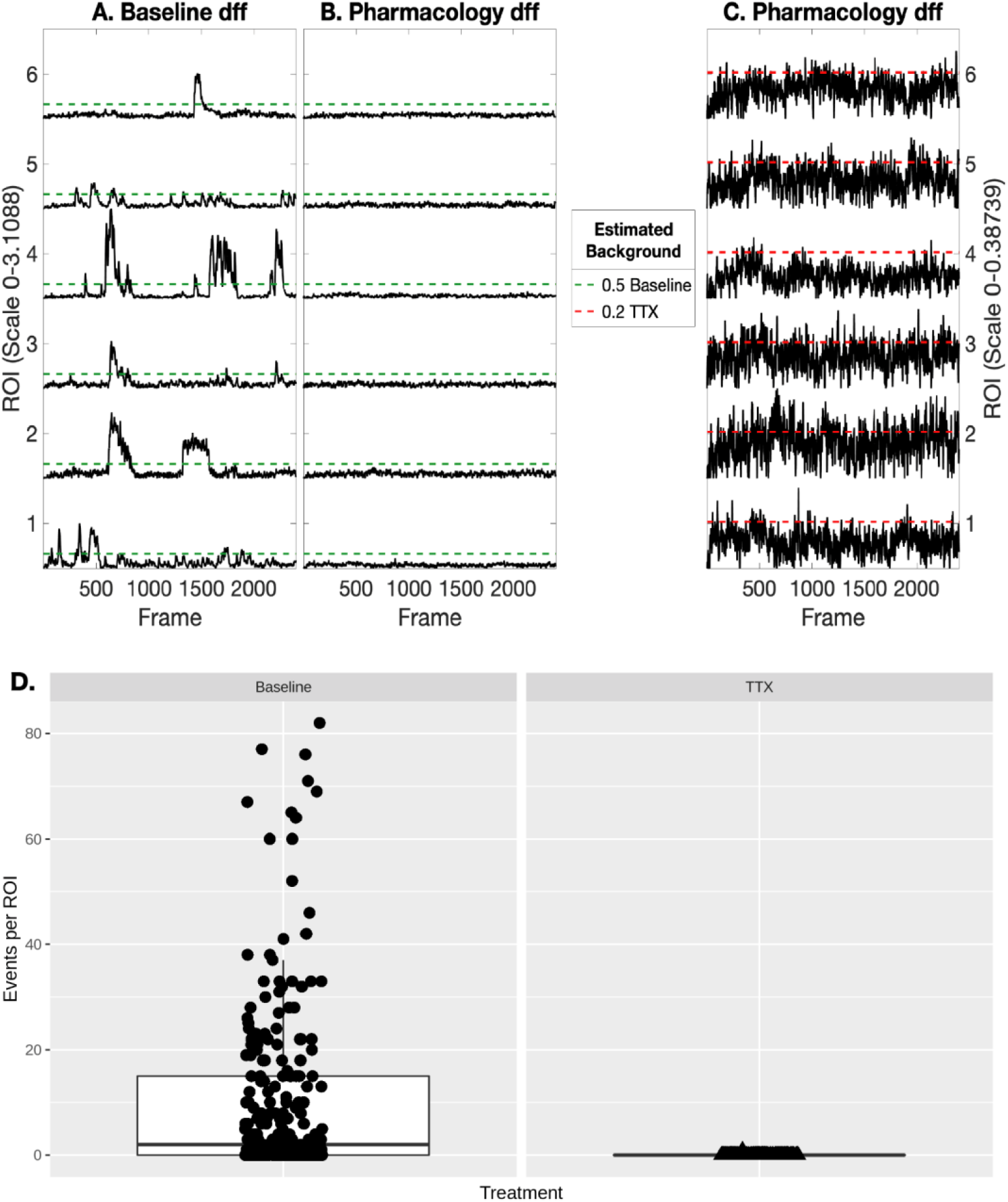
Pharmacological blockade of synaptic transmission illustrates the specificity of CaPTure: **A**) Graph showing normalized calcium activity from a sample baseline field and the estimated background (0.5). **B**) Graph showing normalized calcium activity of the same field treated with TTX, with thresholds from baseline activity to eliminate background. **C**) Graph showing normalized calcium activity of the same pharmacology sample from (B) with the respective intensity scale (y-axis) and background estimated from the same. **D**) Boxplots showing the calcium activity of neurons from baseline and TTX treated fields.

When all data from a given dataset was processed, we compiled all metrics (Step 7). We used the extracted metrics to make comparisons across different experimental manipulations, and made a custom R script for further analysis, to compare the frequency and type of events between neurons derived from individuals diagnosed with schizophrenia and neurotypical controls as previously described [14].

## CONCLUSIONS

Here we have demonstrated the utility of CaPTure to segment neurons and to detect and classify calcium events. CaPTure’s advantages include its ability to effectively segment neurons from surrounding neuropil, which can cause noise in the activity traces and reduce the amplitude and prominence of true events. Additionally, CaPTure uses intensity normalization (df/f) to remove background noise, resulting in reduced incidence of false positives in the final data. The workflow allows for parallel processing of data from large studies, without requiring significant user input or parameterizations. The motif-based method for picking events gives users more insight about the data, including the shape and duration of events. Additionally, the acquisition of high resolution images of cultured neurons could allow users to perform machine learning-based classification on neurons or traces. Calcium events are considered a proxy for neuronal activity, and thus CaPTure provides a powerful tool for researchers to make assessments about the relative cellular and ensemble activity of neurons in culture.

## AVAILABILITY AND REQUIREMENTS

Project name: CaPTure

Project home page: https://github.com/LieberInstitute/CaPTure

Operating system(s): MAC, Windows, LINUX

Programming language: MATLAB

Other requirements:

1. MATLAB image processing toolbox,
2. MATLAB version 2018a or later,
3. Minimum 8GB RAM.

License: General Public License

Any restrictions to use by non-academics

## ABBREVIATIONS

hiPSC: human induced Pluripotent Stem Cell
GECI: Genetically Encoded Calcium Indicator
ROI: Region of Interest
TTX: tetrodotoxin
DFF, df/f: delta fluorescence/fluorescence

---

**Figure S1.**
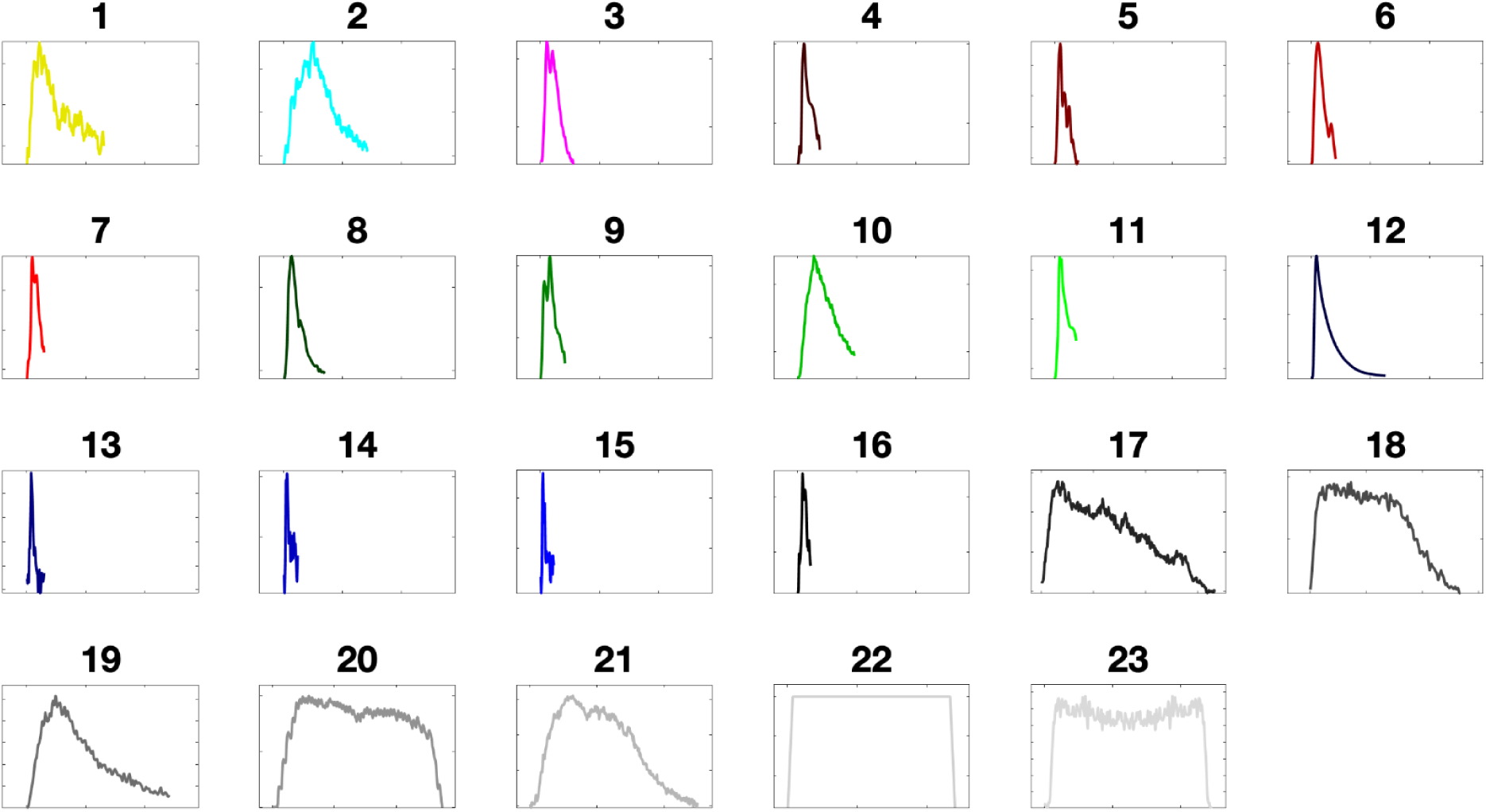
Motif shapes: These plots show the shapes of 23 motifs used in the CaPTure workflow. Motifs 1-16 are adapted from the FluoroSNNAP software and motifs 17-23 were generated based on our data.

## Ethics Approval and Consent to Participate

All human fibroblast donors were part of the Sibling Study of Schizophrenia at the National Institutes of Mental Health in the Clinical Brain Disorders Branch (NIMH, protocol 95M0150, NCT00001486, Annual Report number: ZIA MH002942053, DRW PI). All fibroblast samples were obtained after informed consent.

## Cell Lines

All cell lines used in this work were generated at the Lieber Institute for Brain Development. Identity has been authenticated by STR testing, and mycoplasma testing was performed monthly.

## Availability of data and materials

The datasets analyzed and code used during the current study are available in the Github repository, https://github.com/LieberInstitute/CaPTure.

## Competing Interests

The authors declare that they have no competing interests.

## Funding

This work was funded by the Lieber Institute for Brain Development.

## Author Contributions

Conceptualization: MT, BJM, KM, AEJ, SCP; Methodology: MT, EAP, BAD, CVN, SCP; Investigation: EAP, BAD, CVN, YW, SRS, SCP; Validation: EAP, BAD, SCP; Formal Analysis: MT, SCP; Software: MT; Writing-Original Draft: MT, SCP; Writing-Review and Editing-EAP, BAD, BJM, KM, AEJ; Visualization: MT; Project Administration and Funding Acquisition: BJM, KM, AEJ; Supervision: BJM, KM, AEJ, SCP.

## Acknowledgements

We are grateful for the generosity of the Lieber and Maltz families for making this work possible. This project was supported by the Lieber Institute for Brain Development. We thank Suhaas Adiraju for testing the code, and Kristen Maynard for helpful comments on the manuscript.

## Notes

https://github.com/LieberInstitute/CaPTure

### Competing Interest Statement

The authors have declared no competing interest.

